# Infection with *Listeria monocytogenes* alters the placental transcriptome and eicosanome

**DOI:** 10.1101/2022.04.14.488381

**Authors:** Kayla N. Conner, Derek Holman, Todd Lydic, Jonathan W. Hardy

## Abstract

**Introduction:** Placental infection and inflammation are risk factors for adverse pregnancy outcomes, including preterm labor. However, the mechanisms underlying these outcomes are poorly understood.

**Methods:** To study this response, we have employed a pregnant mouse model of placental infection caused by the bacterial pathogen *Listeria monocyogenes*, which infects the human placenta. Through *in vivo* bioluminescence imaging, we confirm the presence of placental infection and quantify relative infection levels. Infected and control placentas were collected on embryonic day 18 for RNA sequencing to evaluate gene expression signatures associated with infection by *Listeria*.

**Results:** We identified an enrichment of genes associated with eicosanoid biosynthesis, suggesting an increase in eicosanoid production in infected tissues. Because of the known importance of eicosanoids in inflammation and timing of labor, we quantified eicosanoid levels in infected and uninfected placentas using semi-targeted mass spectrometry. We found a significant increase in the concentrations of several key eicosanoids: leukotriene B4, lipoxin A4, prostaglandin A2, prostaglandin D2, and eicosatrienoic acid.

**Discussion:** Our study provides a likely explanation for dysregulation of the timing of labor following placental infection. Further, our results suggest potential biomarkers of placental pathology and targets for clinical intervention.

## Introduction

To ensure the development of the allogeneic fetus, placental immune responses must be precisely balanced between protective immunity and deleterious inflammation [1,2]. Bacterial infection of the placenta can affect this balance, leading to adverse pregnancy outcomes even in the absence of severe disease [1,2]. One such infection is prenatal listeriosis caused by the Gram-positive bacterium *Listeria monocytogenes* (*Lm*). *Lm* is an opportunistic foodborne pathogen that primarily affects the immunocompromised, especially pregnant individuals, who are typically exposed to *Lm* through contaminated meat and dairy products [3]. Following ingestion, *Lm* invades the gut epithelium and traffics in maternal monocytes to the female reproductive organs where it uses cell to cell spread to invade the placenta [3]. Invasion of the placenta can result in a myriad of adverse pregnancy outcomes including preterm labor and downstream abnormal development of the offspring [4–6]. Despite great strides that have been made in the understanding of *Lm* invasion of the placenta, little information is available on the molecular mechanisms underlying listeriosis-associated preterm labor.

Labor and parturition are complicated processes controlled by many genetic, metabolic, and physical factors within the female reproductive tract. Eicosanoids, a family of hormone-like fatty acids, dictate the timing of labor by signaling cervical ripening, breaking down fetal membranes, and promoting myometrial contractility [7–9]. These lipids are produced enzymatically by all cells in the body beginning with the liberation of arachidonic acid from cell membrane phospholipids [10]. Downstream processing by cyclooxygenase (COX) and lipoxygenase (LOX) enzymes yields the two eicosanoid classes: prostaglandins and lipoxins, respectively [10]. Eicosanoids are key players in the delicate balance between protective immunity and deleterious inflammation throughout the body, including the placenta [10]. While associations have been made between eicosanoid pathway perturbations and placental pathology, little information exists regarding infection-induced perturbations to the eicosanoid pathway and downstream consequences in the placenta.

Due to its well characterized lifecycle and genetic malleability, *Lm* has been used as a model for placental infection for decades [11]. In this study, we use a pregnant CD1 mouse model of bioluminescent *Lm* placental infection to begin exploring infection-induced eicosanoid pathway perturbations. We demonstrate through RNA sequencing that mouse placentas colonized with *Lm* have gene expression profiles associated with placental dysfunction and preterm labor. We verify, using semi-targeted mass spectrometry, that these aberrant gene expression profiles result in significant changes to placental eicosanoid concentrations, which we refer to as the placental eicosanome. Together, our data identify a likely mechanism for the induction of preterm labor associated with placental listeriosis infection.

## Materials and Methods

### Strains/Bacterial Culture

The bacterial strain used in this study is the bioluminescent *Listeria monocytogenes* strain Xen32 (Perkin Elmer, Inc.). Cultures were grown overnight, shaking at 37°C in brain heart infusion (BHI) broth supplemented with kanamycin for selection. On the day of mouse infection, overnight cultures were subcultured in fresh BHI supplemented with kanamycin for selection and grown to an OD_600_ of 0.5. The subculture was then diluted in sterile phosphate buffered saline (PBS) to yield 10^6^ colony forming units (CFU) per mL.

### Animals and *In Vivo* Imaging

All mouse experiments were approved by the Institutional Animal Care and Use Committees at Michigan State University and Stanford University. Mice were housed at the Stanford University Research Animal Facility and the Michigan State University Clinical Center animal facility under the care of Campus Animal Resources. The BSL-2 animal procedures were approved under Stanford University Protocol 12342 (formerly 8158) and Michigan State University Animal Use Protocol 201800030. Timed gestation day 11 (E11) pregnant CD-1 mice were delivered on that day from Charles River Laboratories. On E14.5, mice were infected via tail vein injection with 2 × 10^5^ CFU of *Listeria monocytogenes* Xen32 in 200μL phosphate buffered saline prepared as described above *(see “Strains/Bacterial Culture*”). Uninfected control mice were not injected. On E18.5, mice were imaged using the PerkinElmer In Vivo Imaging System (IVIS) to confirm placental infection, then humanely sacrificed under anesthesia according to approved guidelines. Uterine horns were immediately excised and imaged separately using the IVIS to identify infected placentas. Placentas were excised and snap frozen on dry ice then frozen at -80°C for downstream analyses. All animals were imaged using the IVIS for 5 minutes prior to euthanasia, and uterine horns were imaged for 1 minute following excision. Image analysis was performed using the Living Image software by Caliper Life Sciences, and average radiance (light intensity) is expressed as photons per second per centimeter squared per steradian (photons/s/cm^2^/str).

### RNA Sequencing

Twenty infected and four uninfected mouse placentas were excised for downstream RNA sequencing (RNAseq). Tissues were snap frozen on dry ice and stored at -80°C until homogenization. Tissues were homogenized by suspending them in Qiagen Buffer RLT and passing them each subsequently through 16G, 18G, 20G, and 22G needles. Total RNA was extracted from each placenta using the Qiagen RNeasy Midi kit and DNase treated with DNase I (Qiagen) according to the manufacturer’s instructions. Isolated RNA was analyzed for RNA integrity (RIN) values by the Stanford PAN Facility prior to submission for RNAseq analysis by SeqMatic Inc., Mountain View, CA. Single-read sequencing on libraries was performed using the Illumina Genome Analyzer IIx. Data was analyzed on the Galaxy webserver [12]. Raw read files from RNAseq analysis were assessed for quality using FastQC [13], and adapters were removed using Trimmomatic sliding window trimming [14]. To align reads to the mouse reference genome (GRCm39), we used Bowtie2 [15], and resulting alignment files were analyzed for read counts with FeatureCounts [16]. Finally, differential expression analysis was carried out using DESeq2 [17]. Gene ontology and pathway analyses were performed by submitting respective lists for significantly up- and down-regulated genes to g:Profiler with default options (https://biit.cs.ut.ee/gprofiler/gost) [18]. Gene ontology networks were generated using GOnet with custom GO terms related to the eicosanoid pathway (https://tools.dice-database.org/GOnet/) [19].

### Lipidomics

Semi-targeted mass spectrometry (MS) analysis was performed on six infected and six uninfected mouse placentas that had been snap frozen and kept at - 80°C. Placentas were homogenized in methanol acidified with formic acid. Samples were then incubated overnight at -20°C for protein precipitation, then centrifuged. Supernatants were subjected to solid phase extraction using Phenomenex Strata-X 33-micron SPE columns as previously described to concentrate eicosanoids and remove biological matrix components. Eluates were reconstituted in methanol containing 0.01% butylated hydroxytoluene, then centrifuged immediately prior to analysis. Fatty acids and their oxygenated derivatives were analyzed by high resolution/accurate mass (HRAM)-LC-MS. Data-dependent product ion spectra were collected on the four most abundant ions at 30,000x resolution using the FT analyzer. Lipidomics data was analyzed using the Metaboanlyst software according to statistical methods previously published by Xi *et al* [20,21]. Lipid concentrations were normalized to placenta mass, log transformed, and subjected to Pareto scaling prior to statistical analyses.

### Data Availability

Raw sequencing files, normalized count tables, and DESeq2 outputs can be accessed through the NCBI Gene Expression Omnibus.

## Results

### Placental infection by *Lm* alters placental gene expression

At the dose of 2×10^5^ colony forming units (CFU) of bioluminescent *Lm* in pregnant CD1 mice, a range of infection levels is observed across placentas in a single uterine horn, which permits the analysis of many outcomes of prenatal listeriosis including stillbirth and fetal abnormality [22]. Using this model, *in vivo* bioluminescence imaging (BLI) was employed to identify and isolate infected placentas (**Fig 1**). RNA sequencing analysis of 20 infected placentas and 4 control placentas from uninfected animals revealed 498 significantly underexpressed and 862 significantly overexpressed (Log_2_FC ≤ -1 or ≥ 1; adjusted P value ≤ 0.05) in the infected placentas **(Fig. 2A, Supplementary Material)**. The top five overexpressed genes following infection included *Zbp1, GM12250, Igtp, Tap1*, and *Ido1* **(Fig. 2A, Supplementary Material)**. These results were expected considering the various immunoregulatory roles these genes are known to play. We observed minimal variability in the four uninfected sample gene expression profiles, which formed their own distinct cluster (Fig. 2B). Conversely, the infected sample gene expression profiles displayed considerable variability, which is consistent with the range of infection levels in our model (**Fig. 1, Fig. 2B**).

**Figure 1.**
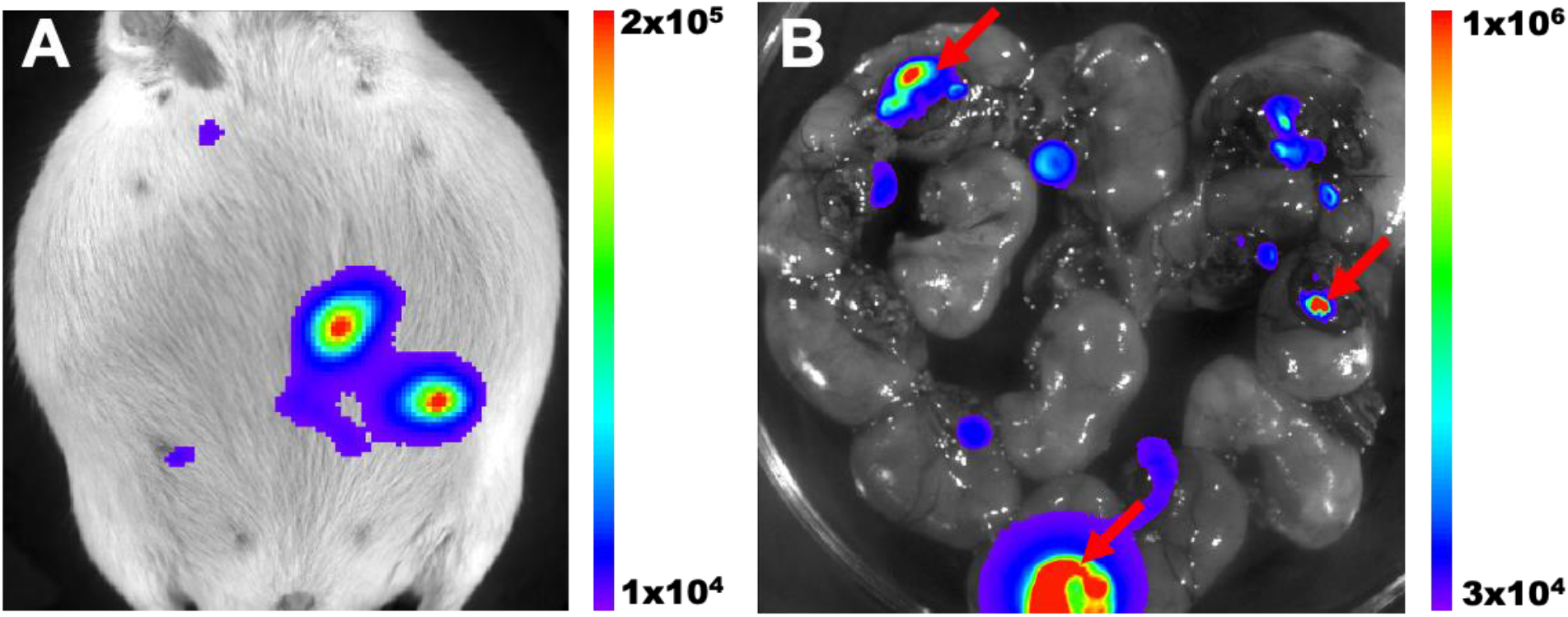
*In vivo* bioluminescence imaging of *Lm* in the placenta. (A) Example of bioluminescence imaging of a pregnant mouse infected with *Lm* on E14.5 and imaged on E18.5. (B) Excised uterine horns from a similar animal showing the placentas used for RNAseq. RNA from infected placentas (arrows) was sequenced and compared to controls from uninfected mice. The false color scale is photons/second.

**Figure 2.**
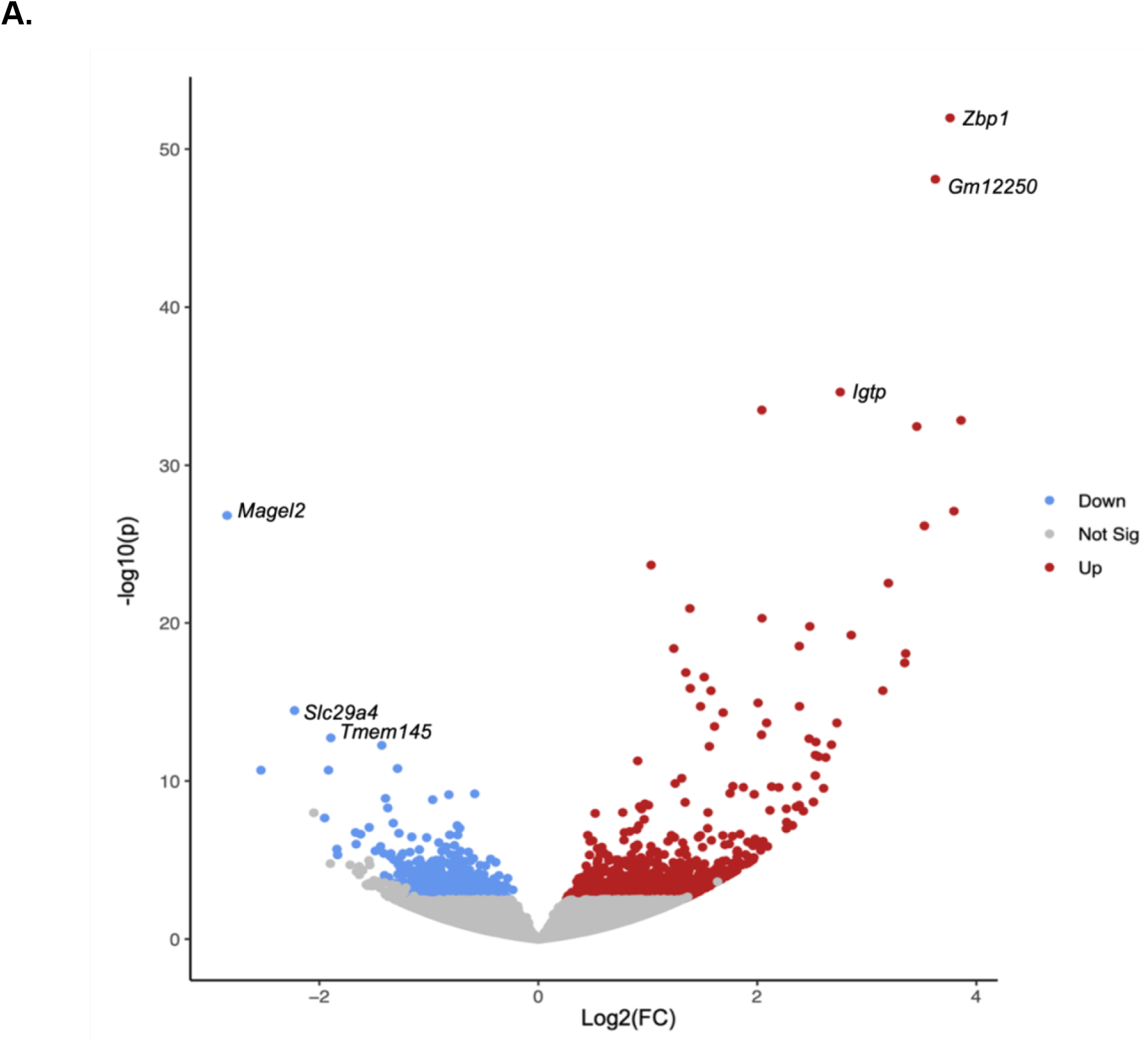
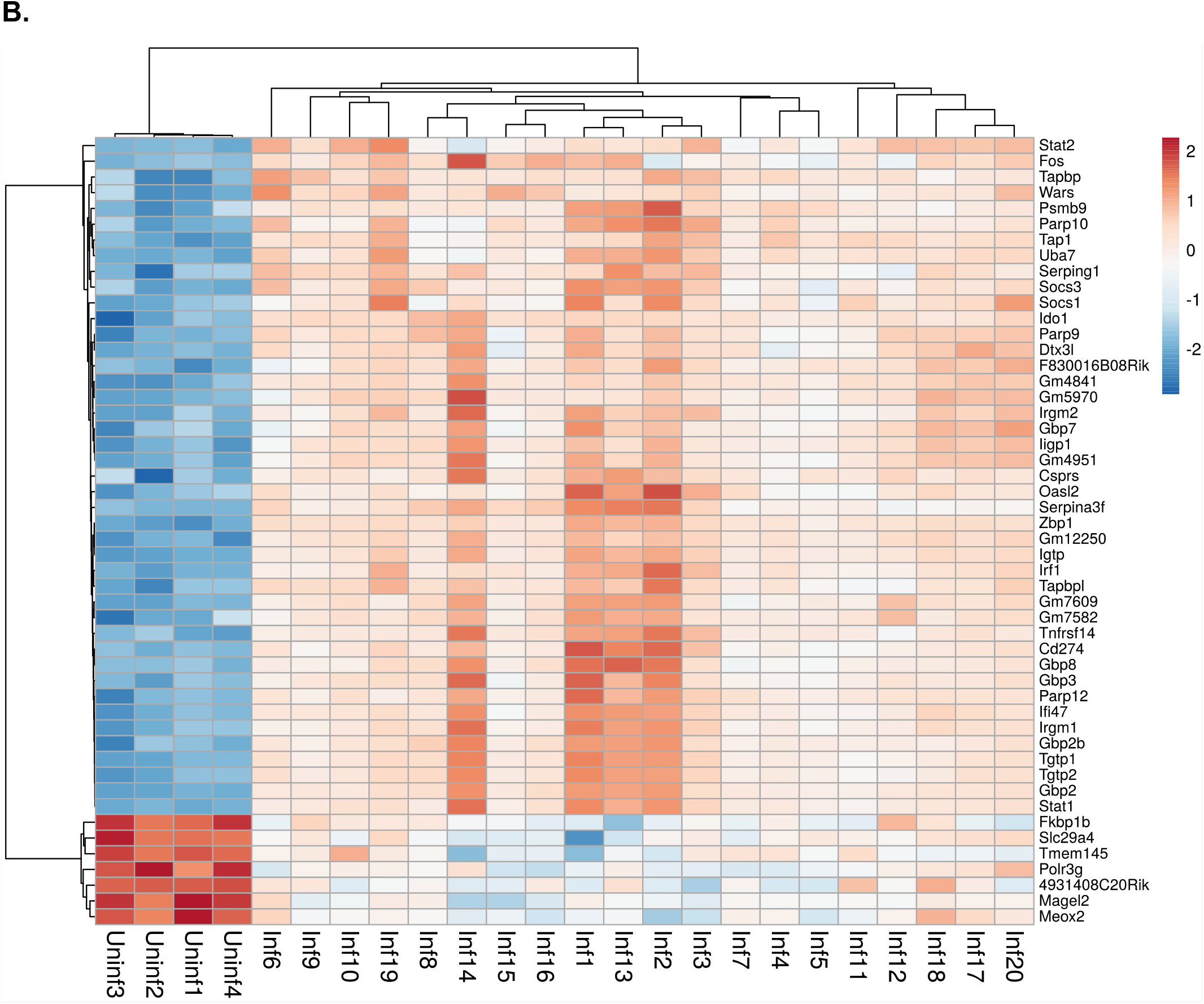
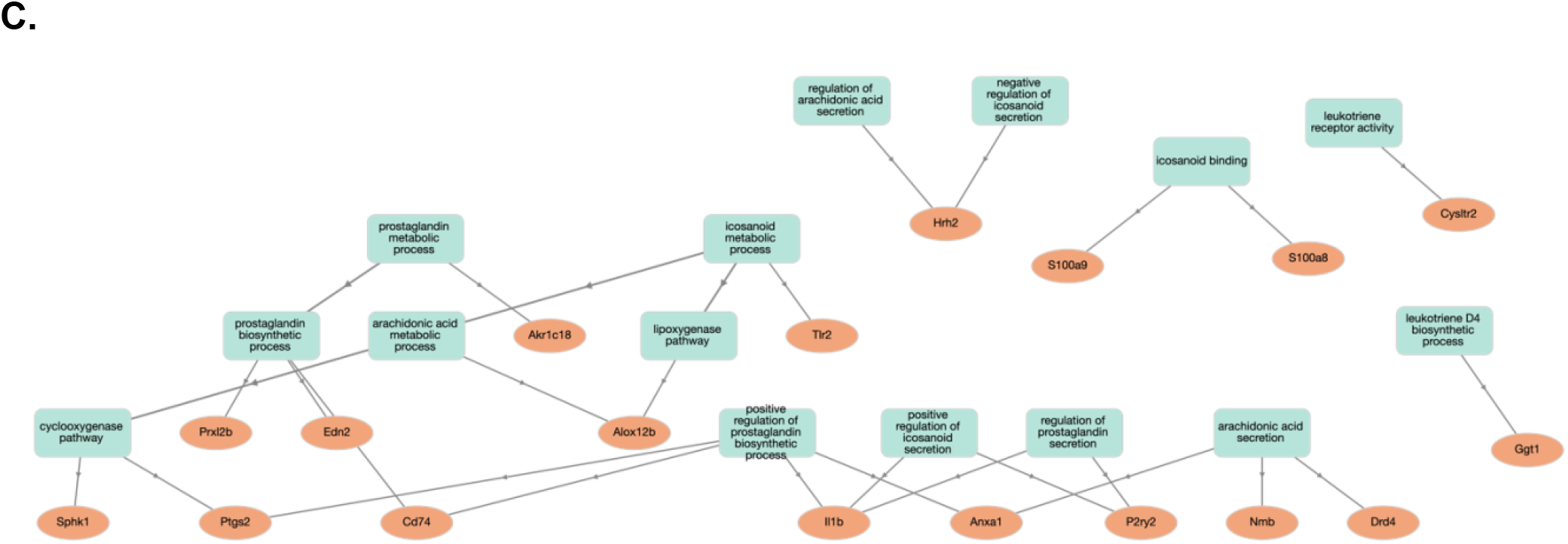
Gene expression profiles are altered in *Lm*-infected placentas. Differentially expressed genes in *Listeria*-infected placentas (compared to uninfected) were determined using DEseq2 and expressed as a volcano plot (A). Significantly overexpressed genes (fold change ≥ 2; Adjusted P value ≤ 0.05) are highlighted in red while significantly underexpressed genes (fold change ≤ -2; Adjusted P value ≤ 0.05) are highlighted in blue. Values are presented as Log_2_ Fold Change and Log_10_ Adjusted P Value. The top 50 differentially expressed genes are expressed as a heatmap of normalized counts per sample (B). Heatmap was generated using Heatmapper [35], and sample clustering was computed with average linkage clustering and Euclidian distance measurement (represented by the sample dendogram). Gene ontology analysis was conducted using g:profiler, and network visualization was generated using GOnet (C). GO terms are in blue-green rectangles and gene names are in orange ovals.

To better understand the pathways associated with significantly dysregulated genes, we performed functional profiling using g:Profiler. This analysis revealed several pathways of interest in both up- and down-regulated gene data sets (**Table S2, Supplementary Material**). Interestingly, pathways associated with underexpressed genes were largely related to ion transport across the membrane (**Supplementary Material**). As expected, most pathways associated with overexpressed genes were linked to pro-inflammatory processes typical of bacterial infection, consistent with the expected infiltration and activation of immune cells (**Supplementary Material**). Notably, GO terms related to prostanoid and prostaglandin biosynthesis were enriched in our upregulated gene data set (**Supplementary Material**). To visualize overexpressed gene networks associated with eicosanoid metabolism, we submitted our overexpressed genes and custom gene ontology ID list to GOnet to generate a visual custom gene ontology network (**Fig. 2C**).

Following gene ontology analysis, we became interested in the enrichment of eicosanoid metabolism genes due to the known roles of eicosanoids in pregnancy and listeriosis elsewhere in the body. In our RNAseq data, we observed a significant overexpression (approximately 2.3-fold increase, adjusted P value ≤ 0.05) in the *Ptgs2* gene encoding cyclooxygenase 2, a key enzyme in the eicosanoid pathway (**Fig. 2C, Supplementary Material**). While *Ptgs2* encoding cyclooxygenase 2 was significantly overexpressed, the *Ptgs1* gene encoding cyclooxygenase 1 (the constitutive housekeeping isoform of this enzyme) was not significantly dysregulated (**Supplementary Material**). In addition to *Ptgs2*, we observed overexpression of several other eicosanoid-associated genes (**Table S1, Fig. 2C**). We hypothesized that, due to overexpression of several genes associated with eicosanoid production, the concentrations of these lipids would be increased in infected placentas. Specifically, we hypothesized that infected placentas would harbor increased concentrations of prostaglandins due to the upregulation of several enzymes implicated in prostaglandin synthesis (**Fig. 3**). In addition, because eicosanoid pathway enzymes can be regulated by post-transcriptional mechanisms including allosteric induction [23], it was important to measure the pathway products themselves to fully characterize changes in this pathway.

**Figure 3.**
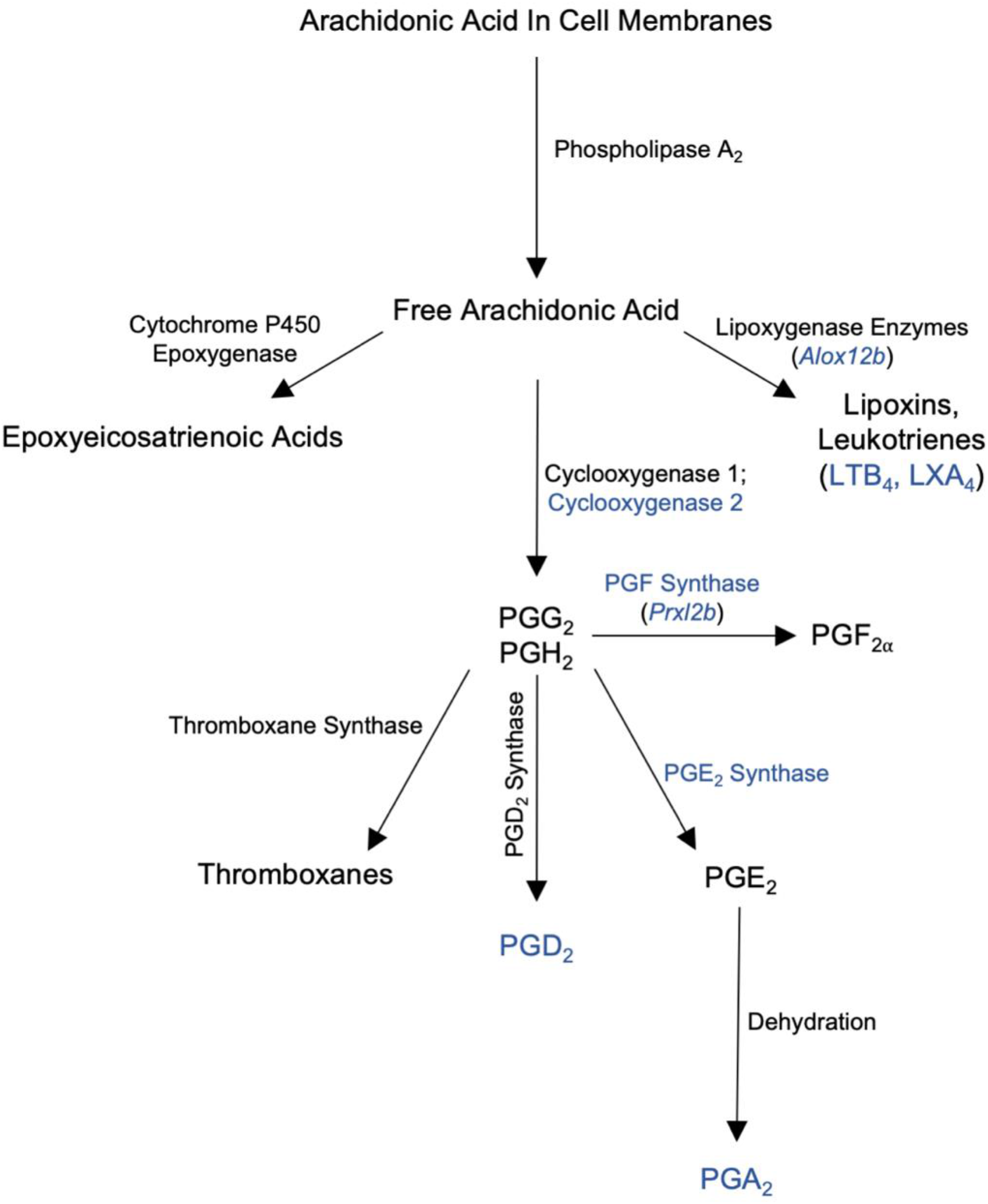
*Lm* infection results in upregulation of key eicosanoid pathway enzymes and increased concentrations of specific eicosanoids in the placenta. This adapted eicosanoid pathway figure illustrates the points at which this pathway is altered by listeriosis in the placenta. Genes for enzymes represented in blue text as well as eicosanoids represented in blue text are significantly overexpressed in our data sets.

### *Lm* infection alters eicosanoid concentrations in the placenta

Because we wanted to know if eicosanoid levels were perturbed along with eicosanoid pathway gene expression, we carried out semi-targeted mass spectrometry to measure concentrations of various eicosanoids in infected and uninfected placentas (**Fig. 1**). Our analysis revealed distinct profiles for infected versus uninfected placentas **(Fig. 4**). We observed 12 eicosanoids showing a ≥2-fold increase or decrease in concentration in the placenta following infection with *Lm* (**Fig. 4**). Strikingly, leukotriene B_4_ (LTB_4_) exhibited a ∼25-fold increase following infection (**Fig. 4, Supplementary Material**). Also of note were prostaglandin A_2_ (PGA2), prostaglandin E_2_ (PGE_2_), prostaglandin D_2_ (PGD_2_), and prostaglandin F_2α_ (PGF_2α_) which showed ∼4.8-, ∼2.4-, ∼2.1, and ∼2.3-fold increases following infection, respectively (**Fig. 4, Supplementary Material**). Of these dysregulated eicosanoids, nine reached statistical significance (p ≤ 0.05) including LTB_4_, LXA_4_, PGA_2_, PGD_2_, and eicosatrienoic acid (**Fig. 5, Supplementary Material** Together, these data supported our hypothesis that altered gene expression in the placenta results in changes in placental eicosanoid profiles, which we refer to as the placental eicosonome.

**Figure 4.**
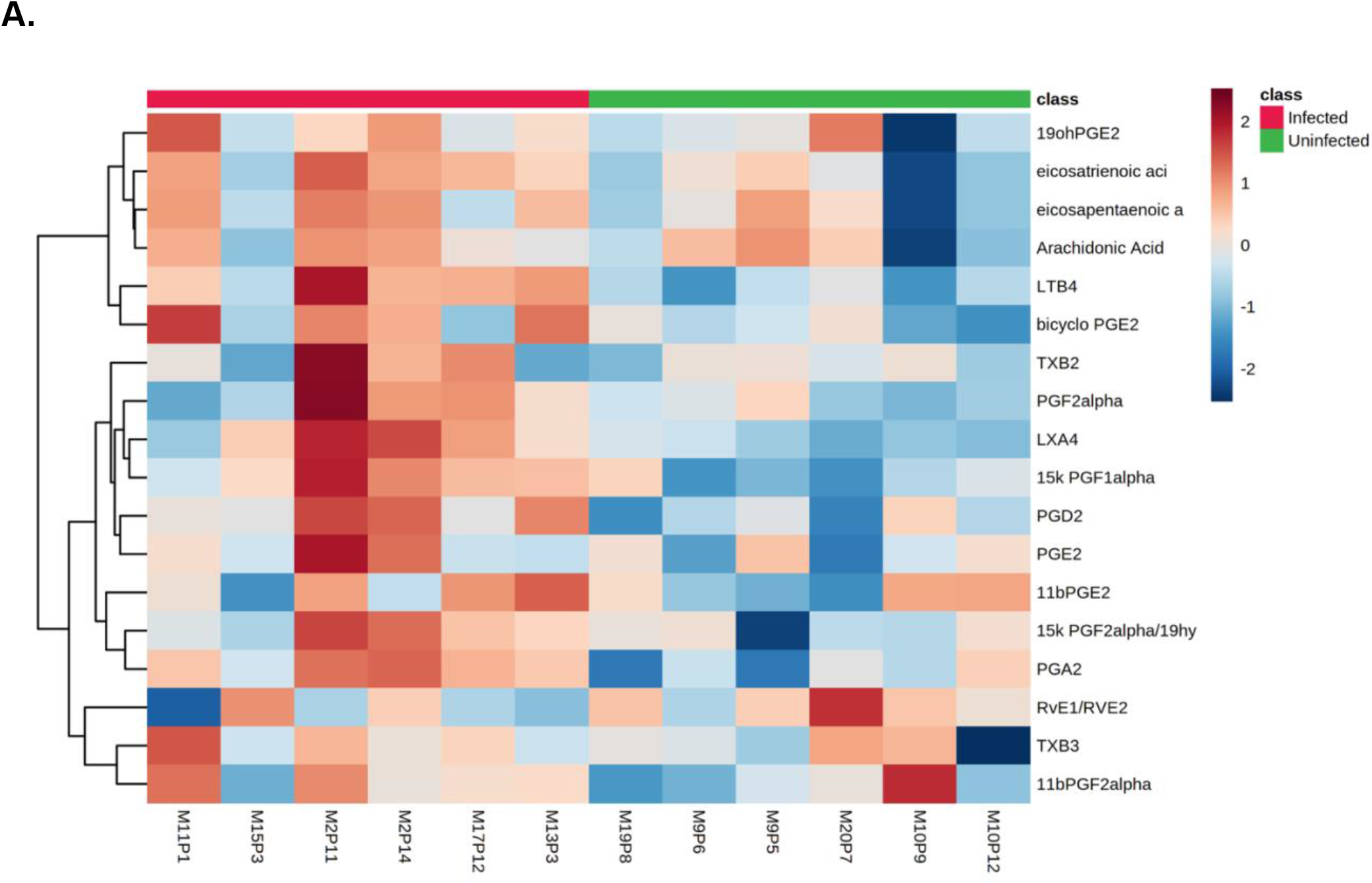
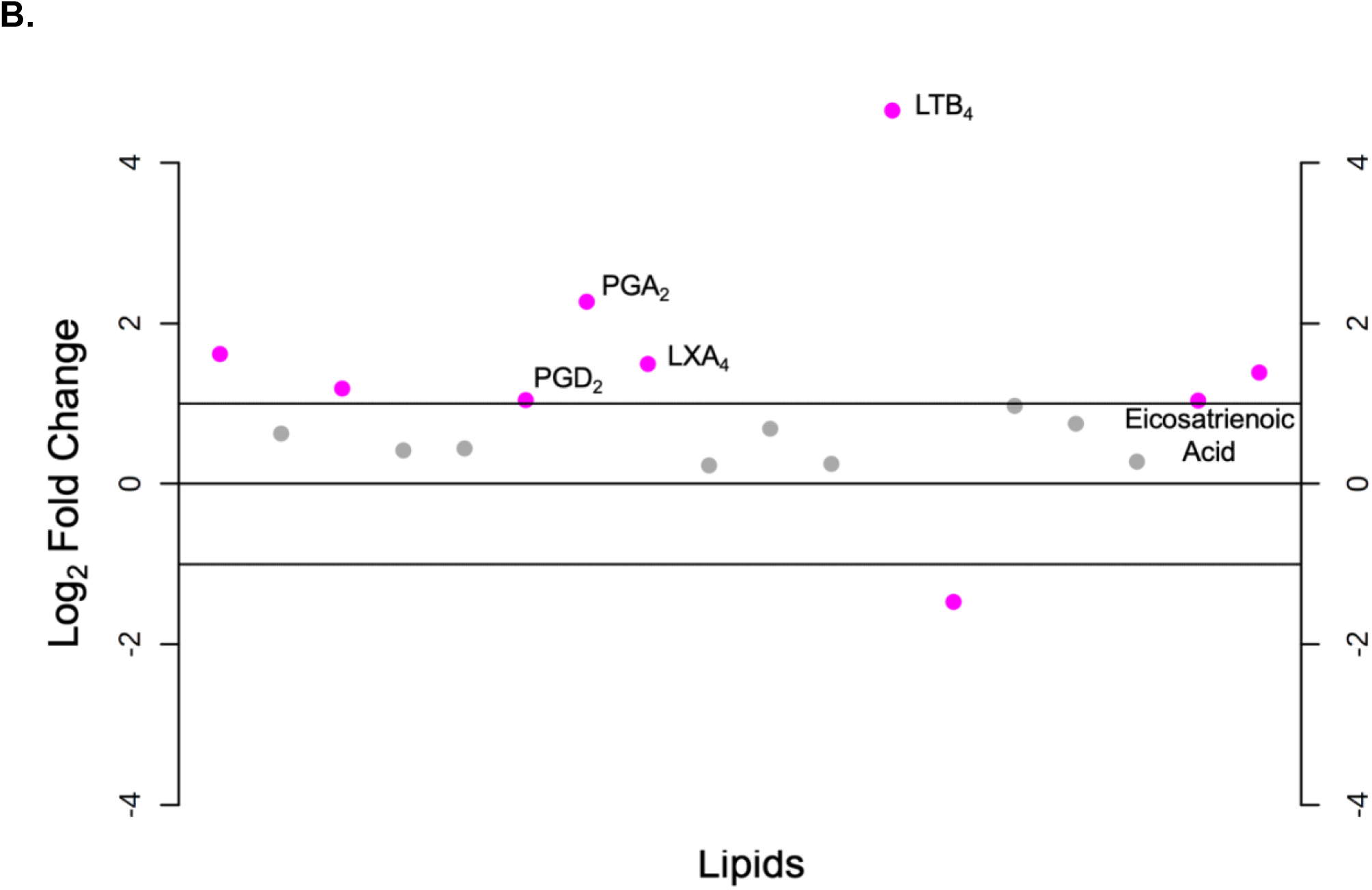
*Lm* infection alters the placental eicosanome. Eicosanoid profiles for infected and uninfected placentas were assessed using semi-targeted mass spectrometry. A heatmap was generated using Metaboanalyst to compare relative eicosanoid concentrations in infected versus uninfected placental samples (A). Fold change was analyzed using Metaboanalyst and is expressed as a dot plot with each dot representing the Log2 fold change (infected/uninfected) of each compound in our eicosanoid panel (B). Eicosanoids with >2-fold change are represented by pink dots, and significantly overexpressed eicosanoids are labeled.

**Figure 5.**
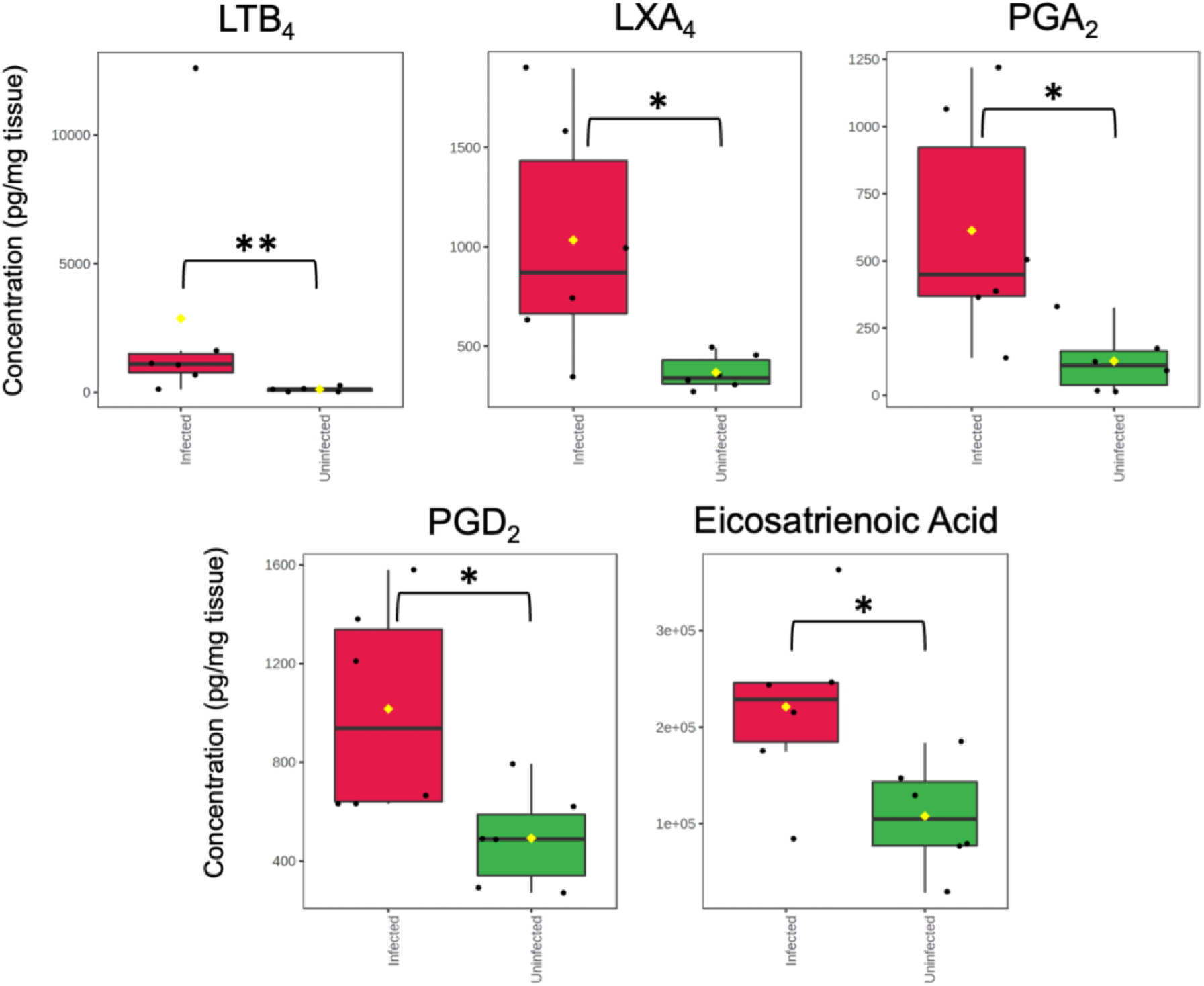
Several eicosanoids are significantly overexpressed following placental infection. Eicosanoid profiles for infected and uninfected placentas were assessed using semi-targeted mass spectrometry. Data was analyzed using Metaboanalyst, and eicosanoids which reached statistical significance (*p < 0.05, **p < 0.01) are represented as box plots below. Infected samples are in red while uninfected samples are in green. Each dot represents one sample. Concentrations are expressed as pg/mg of tissue.

## Discussion

Pregnancy complications including preterm birth are relatively common, and preterm birth is the leading cause of infant mortality worldwide [24,25]. While many factors can contribute to the occurrence of preterm birth, the outcome can be developmentally devastating for the infant. Infants born prematurely are more likely to exhibit breathing problems, sensory problems, and developmental delay [26]. Infection is a well-known cause of preterm birth, necessitating studies of prenatal responses to distinct pathogens [25]. Because of the crucial role of the placenta in immune responses during pregnancy, pathogens that infect this organ are especially important to understand. For example, it will be crucial to distinguish placental infection from other prenatal infections such as chorioamnionitis, which may elicit completely different responses and require different interventions. In addition, placental infection can induce inflammatory responses, which have been associated with preterm birth [27]. Therefore, animal models of placental infection are vital tools in understanding preterm birth.

*Listeria monocytogenes* is a known placental pathogen that can cause preterm labor as well as other perinatal pathologies [11,28]. Animal models of prenatal listeriosis have revealed details of placental infection, including the target cell type, bacterial virulence factors and molecular mechanisms of invasion [29]. However, host placental responses to this bacterium have not been previously defined and may reveal clues as to the function of the placenta in prenatal resistance to infection. Our data sheds light on the molecular and metabolic mechanisms underlying listeriosis-induced preterm labor. We have shown using a pregnant mouse model of placental listeriosis that infected placentas harbor distinct gene expression profiles compared to their uninfected counterparts. Unsurprisingly, we have identified an enrichment of genes associated with inflammation and response to infection in infected placentas. We were particularly interested to observe an enrichment of genes associated with eicosanoid biosynthesis and metabolism following infection. Though this result is not entirely surprising due to the role of eicosanoids in inflammation, it was noteworthy considering that eicosanoids are known to play critical roles in the regulation of labor, as well as other aspects of pregnancy such as placental function [30]. Further, this discovery warrants further investigation due to previous associations between placental eicosanoid dysregulation and pathological pregnancy outcomes in previous studies [31].

To determine if eicosanoid concentrations were perturbed along with gene expression profiles, we employed a semi-targeted mass spectrometry approach to quantify the eicosanoid concentrations in infected and uninfected mouse placentas. This analysis highlighted perturbations in eicosanoid concentrations in infected placentas. We noted significant increases in the concentrations of LTB_4_, LXA_4_, PGA_2_, PGD_2_, and eicosatrienoic acid. Previous studies strongly support the association between the eicosanoids we have identified as increased in placental infection and placental pathology, including LTB_4_ [32], LXA_4_ [33], and PGD_2_ [34].

Broadening our understanding of molecular mechanisms underlying listeriosis-induced adverse pregnancy outcomes has the potential to propel the development of improved clinical interventions for pregnancy associated listeriosis and other placental infections. Our study offers insight into the genetic and metabolic changes that take place in the placenta following *Lm* infection. While our study begins to offer possible mechanisms of listeriosis-induced preterm labor, much remains to be investigated.

To our knowledge, this is the first study associating increased PGA_2_ concentrations with placental infection or preterm labor. This is noteworthy as PGA_2_ is a known degradation product resulting from the dehydration of PGE_2_, which has been studied extensively for its role in the timing and induction of parturition. Increased PGA_2_ concentrations could imply an increase in upstream PGE_2_ production and its subsequent degradation, which could be contribute to dysregulation of labor. Future studies should address the mechanistic role of this eicosanoid in the context of infection-induced preterm labor.

Our observations confirm that the known role of eicosanoids in infection and inflammation in other tissues also applies to the placenta, where the eicosanoids are also known to function in the timing of labor. It is noteworthy that many of the eicosanoids identified in our study have been implicated in pathological pregnancy outcomes and placental disease. In addition, the induction of specific prostaglandins and leukotrienes suggests the possibility of receptor-specific interventions. It is important to identify new detection and intervention methods that can be utilized to prevent adverse pregnancy outcome. We propose that future studies assess eicosanoid concentrations in maternal circulation to assess the usefulness of eicosanoids as clinical biomarkers of placental disease. Further, we suggest that eicosanoid synthesis and uptake be studied as a potential route of intervention in the prevention of infection-induced adverse pregnancy outcome.

## Supporting information

Supplemental File - DESeq2

Supplemental File - Gene Ontology Underexpressed

Supplemental File - Gene Ontology Overexpressed

## Author Contributions

Conceptualization: JWH, KNC; Investigation: KNC, DH, TL, JWH; Analysis: TL, KNC; Writing: All Authors; Visualization: KNC; Project Administration: JWH; Resources and Funding Acquisition: JWH.

